# Ecological fNIRS in mobile children: Using short separation channels to correct for systemic contamination during naturalistic neuroimaging

**DOI:** 10.1101/2024.06.02.596560

**Authors:** P. Pinti, L.M Dina, T Smith

**Affiliations:** Department of Psychological Sciences, Birkbeck, University of London WC1E 7JL, United Kingdom; Department of Medical Physics and Biomedical Engineering, University College London, United Kingdom; Department of Psychology, King’s College London, London SE5 8AB, United Kingdom; Creative Computing Institute, University of the Arts London SE5 8UF, United Kingdom

**Keywords:** short separation channels, fNIRS, superficial signal regression, children, freely moving, naturalistic, systemic interferences

## Abstract

**Significance:** The advances and the miniaturization in functional Near Infrared Spectroscopy (fNIRS) instrumentation offers the potential to move the classical laboratory-based cognitive neuroscience investigations into more naturalistic settings. Wearable and mobile fNIRS devices also provide a novel child-friendly means to image functional brain activity in freely moving toddlers and preschoolers. Measuring brain activity in more ecologically valid settings with fNIRS presents additional challenges, such as the increased impact of physiological interferences. One of the most popular methods to minimize such interferences is to regress out short separation channels from the long separation channels (i.e., superficial signal regression or SSR). Whilst this has been extensively investigated in adults, little is known about the impact of systemic changes on the fNIRS signals recorded in children in either classical or novel naturalistic experiments.

**Aim:** We aim to investigate if extracerebral physiological changes occur in toddlers and preschoolers, and whether SSR can help minimize these interferences.

**Approach:** We collected fNIRS data from 3-to-7 years olds during a conventional computerized static task and in a dynamic naturalistic task in an immersive virtual reality (VR) continuous automatic virtual environment (CAVE).

**Results:** Our results show that superficial signal contamination data is present in both young children as in adults. Importantly, we find that SSR helps in improving the localization of functional brain activity, both in the computerized task and, to a larger extent, in the dynamic VR task.

**Conclusions:** Following from these results, we formulate suggestions to advance the field of developmental neuroimaging with fNIRS, particularly in ecological settings.

## Introduction

Functional Near Infrared Spectroscopy (fNIRS) is a non-invasive neuroimaging modality that measures the cortical haemodynamic and oxygenation changes that follow neuronal activity. Thanks to its being portable, relatively robust to head movements, versatile for a wide range of population and tasks, fNIRS has gained increased popularity over the past 30 years, as recently acknowledged in the “Celebrating the 30 years of fNIRS” special issue in Neurophotonics (Highton et al., 2023). The tremendous progress in hardware developments has led to the availability of wireless/mobile fNIRS devices that now enable brain function imaging in real-world settings and in those situations where other neuroimaging methods like functional magnetic resonance imaging (fMRI) or electroencephalography (EEG) are not suitable, such as those requiring dynamic movements (Balardin et al., 2017; Burgess et al., 2022; Greaves et al., 2022) or face-to-face social interactions (Krishnan-Barman et al., 2023; Nguyen et al., 2021).

fNIRS is very often used within the field of cognitive neuroscience to localize the task-evoked functional brain activity or to evaluate the brain-to-brain or within brain functional connectivity (Yücel et al., 2021). Results are typically assessed at the group level rather than at the individual level by testing specific hypotheses on the pooled single subjects’ data. The reliability and robustness of the inference results strongly rely on the amount of noise in the included fNIRS data. In fact, the fNIRS signals are contaminated by noise components of different origin: measurement noise (e.g., electronic noise), motion artifacts, and physiological noise (e.g., changes in blood pressure, respiration rate, heart rate (Pinti et al., 2019; Yücel et al., 2021)). These can act as confounding factors in the fNIRS analysis and can either mask and/or mimic the presence of a task-evoked hemodynamic response, leading to false positive and/or false negatives at the group level statistics (Tachtsidis & Scholkmann, 2016). Some of these components have distinct frequency characteristics from the task-evoked hemodynamic response and hence can be easily removed or minimized using filtering methods (e.g., band-pass filter to remove very low and high frequency noise, or wavelet filtering to correct for motion artifacts (Scholkmann et al., 2014)). However, other physiological confounders can overlap with the task-evoked hemodynamic activity, such as variations in arterial blood pressure known as Mayer waves (∼0.1 Hz (Yücel et al., 2016)) or respiration (∼0.2 Hz), and more advanced methods are needed to minimize their impact.

Among these, one major source of interference on the cortical fNIRS signals comes from the extra-cerebral contamination: the near infrared light emitted from the light source travels twice through various layers of the head (skin, skull, dura, cerebrospinal fluid) before reaching the brain and being back-scattered to the detector (Tachtsidis & Scholkmann, 2016). Previous work has shown that approximately 96% of the injected light is absorbed in the skin and skull, while only 3% is absorbed in the brain (Haeussinger et al., 2011). Therefore, the measured fNIRS signal is a mixture of the hemodynamic and oxygenation changes happening in both the superficial *and* cortical vasculature. Additionally, blood flow changes in both the brain and the scalp can be modulated by those physiological processes that act on the vascular tone. These processes encompass changes in blood pressure, respiration, partial pressure of CO_2_ (PaCO_2_), and autonomic activity, among others. These elements can either be spontaneous (i.e., heartbeat, respiration, variations in arterial blood pressure or Mayer waves, very low frequencies) or modulated or induced by the task itself. For example, posture changes can alter blood pressure (Tachtsidis et al., 2004), speaking and jaw movements can lead to changes in PaCO_2_ and non-neural blood flow changes related to the use of the temporalis muscle (Zimeo Morais et al., 2017), as well as stressful tasks can promote vasoconstriction by triggering a sympathetic response (Pinti et al., 2015). Even passive and apparently stress-free tasks such as passive color light exposure can induce physiological changes (Scholkmann et al., 2017). It is thus reasonable to expect even a larger impact of systemic interferences when fNIRS neuroimaging experiments are extended from the stationary laboratory setting to those with a higher degree of ecological validity where participants may be able to walk, engage in dynamic movements or speak freely (Pinti et al., 2018).

Different strategies have been proposed so far to deal with physiological interferences, such as multimodal monitoring or appropriate task design (e.g., avoid 10s-long task and rest blocks that overlaps with Mayer waves) (Yücel et al., 2021), include experimental conditions with the same level of physical activity so some of the effects can be subtracted when contrasting the conditions (see Burgess et al. (2022) (Burgess et al., 2022) for an example)). A popular method to account for superficial contamination is to use short separation channels alongside long separation channels. In contrast with long separation channel where a pair of source and detector that are placed at a distance >2 cm (typically 2 cm for infants and 3 cm for adults) and are assumed to be sampling from the brain, short separation channels are created by placing a source and a detector at <1 cm distance and assume that the back-scattered photons have travelled through the superficial layers of the head only and are not brain sensitive (Brigadoi & Cooper, 2015). The optical signal from the short separation channel can be regressed out from the long separation channel to obtain a more brain-specific fNIRS signal with reduced scalp contamination. This method is often referred to as *superficial signal regression* (SSR; (Gregg, 2010)). SSR has been widely used in the fNIRS field and has been demonstrated to improve the recovery of the hemodynamic response (Noah at al., 2021), even in instances when strong Mayer waves components are present (Yücel et al., 2016). It was also proven to be effective in increasing the performance of resting-state functional connectivity analyses (Paranawithana et al., 2022).

Whilst the benefit of SSR has been well established in adult populations and nowadays can be considered common practice (Yücel et al., 2021), less is known on its impact on the group level statistics of developmental fNIRS data. Furthermore, it is not routinely used in children’s fNIRS data analysis. It is reasonable to hypothesize that the scalp haemodynamic changes might be different in younger populations compared to adults because of a different head anatomical structure or maturation stage of the vasculature system (Emberson et al., 2016); therefore, it is not clear yet whether there are significant superficial changes in children and whether regressing out the short separation channels can reduce physiological interference and increase the reliability of the task-evoked group level inferences. To date, only the study by Ferradal et al. (2016) (Ferradal et al., 2016) performed SSR in high-density fNIRS data recorded on newborns. However, others have reported task-evoked patterns in channels with a source-detector separation of ∼10 mm in 4-to-7 months old infants that suggest that a proportion of this signal is of superficial origin (Emberson et al., 2016; Frijia et al., 2021). Emberson and colleagues (2016) were the first to investigate whether SSR has an impact on the group level statistics in a developmental sample; however, this was only investigated in infants’ fNIRS data. They showed that both superficial and deeper haemodynamic responses can be observed in the occipital cortex of 6 months old babies while undergoing a visual and auditory stimulation task. However, the removal of superficial signals did not lead to significant changes on the group level results. As pointed out by the authors, these results might be task-specific, region-specific and population-specific, and a different outcome may be expected in other experiments or populations and further investigations are needed.

There remains a notable gap of knowledge regarding the benefits of SSR across development, such as in toddlers and preschoolers. Previous studies have shown that cerebral blood flow is lower in the postnatal brain than in adults, but then increases until 7 years of age and then decreases to a similar level to adults in teenage years, which may reflect brain maturation and synapses formation (Kozberg & Hillman, 2016). Therefore, children older than 3 years old may exhibit a different pattern of blood flow changes which may significantly differ from those of younger or older population. More importantly, task-evoked haemodynamic changes in these age groups may be modulated by experimental contexts (e.g., standing or sitting) and physical movements as previously shown in adults (Pinti et al., 2018; Tachtsidis et al., 2004). With the recent advances in wireless and more mobile NIRS instrumentation, cognitive neuroscience investigations can now move to more naturalistic settings (Vidal-Rosas et al., 2023) which are more dynamic and may thus enhance the impact of physiological interferences on the fNIRS-derived brain signals. Increasing the ecological-validity of fNIRS procedures is of critical importance for studying toddlers and pre-schoolers whose neuro-cognitive development is traditionally under researched due to their difficulty sitting still and complying with task demands.

In this work, we aim to fill this gap and investigate if: 1) there are significant extracerebral physiological and hemodynamic changes in toddlers and preschoolers; 2) extracerebral interference is stronger when fNIRS data are recorded in standing and freely moving children; 3) SSR can mitigate the impact of systemic confounding factors and improve the robustness of fNIRS data; and 4) SSR has a significant effect on the group level inference results on the task-evoked brain activity. To this goal, we collected fNIRS data on a group of 3-to-7 years old children undergoing an inhibitory control task, a core executive function (Diamond, 2013).Participants performed a standard computer version of the task sitting at a desk and a naturalistic virtual reality (VR) version adapted after the computer task. The VR task was carried out in an immersive VR continuous automatic virtual environment (CAVE) where kids could stand and were able to move about. The task was designed to account for possible systemic interferences. Finally, we provide some recommendations on how to adapt some of the best practices (Yücel et al., 2021) to help enable fNIRS neuroimaging on young children.

Our hypotheses are: 1) there are task-evoked changes in scalp blood flow in toddlers and preschoolers; 2) these are larger in the VR version of the task compared to the computer-based one because of posture (standing vs sitting) and physical activity; 3) SSR changes the outcome of the group level statistics especially in the VR task; 4) SSR has a larger effect on oxygenated haemoglobin than deoxygenated haemoglobin as previously found in adults (Kirilina et al., 2012; Tachtsidis & Scholkmann, 2016).

## Material and methods

### Participants

Thirty-nine 3-to-7 years old children (range: 3-7 years, M_age_=4.45, SD=1.08, 35.9% female) were recruited. All participants were born full-term, healthy, with normal or corrected to normal vision and hearing, and with no diagnosis of a neurodevelopmental condition. The protocol for this study was pre-registered on the Open Science Framework (Dinu et al., 2022). All parents or caregivers provided written informed consent. Ethical approval was granted by the Ethics Committee of the Department of Psychological Sciences at Birkbeck, University of London (No. 2021072).

For the VR task, 7 participants were excluded for poor fNIRS data and 2 for task performance; for the computer task, 2 participants were excluded due to poor fNIRS data and performance, 6 for poor fNIRS data only, and one for task performance only. The final sample thus included 30 participants for the VR task (Mage = 4.5, SD = 1.14, 21 males) and 30 participants for the computer task (Mage = 4.53, SD = 1.14, 21 males).

### Experimental protocol

Participants performed a response inhibition task, a type of inhibitory control referring to our ability to suppress a certain action or a prepotent response. The Go/No-Go task is a widely used measure of response inhibition, which require participants to press a button in response to some stimuli and to refrain from button presses when certain other stimuli appear (Diamond, 2013). Here, we used child-friendly versions of a Go/No-Go task (Schröer et al., 2021). In particular, a block-designed Go/No-Go task was performed in two versions: a computerized and a VR version.

*Computer-based Go/No-Go Task (CB).* This version (Figure 1 A) was previously used in Schröer et al. (2021) and restructured as a block-design to fit with fNIRS requirements. Children were sitting in front of a computer screen and were told a story that a town was hunted by vampires and that they had to be monster hunters and catch all the bats that would soon become vampires. They were presented with pictures of bats (Go trials) and cats (No-Go trials). They were asked to press the space bar on the computer keyboard as fast as possible to catch the bats (Go) but refrain from doing so when they would see a cat (No-Go). The task started with two practice trials before moving on to the main experiment.

**Figure 1.**
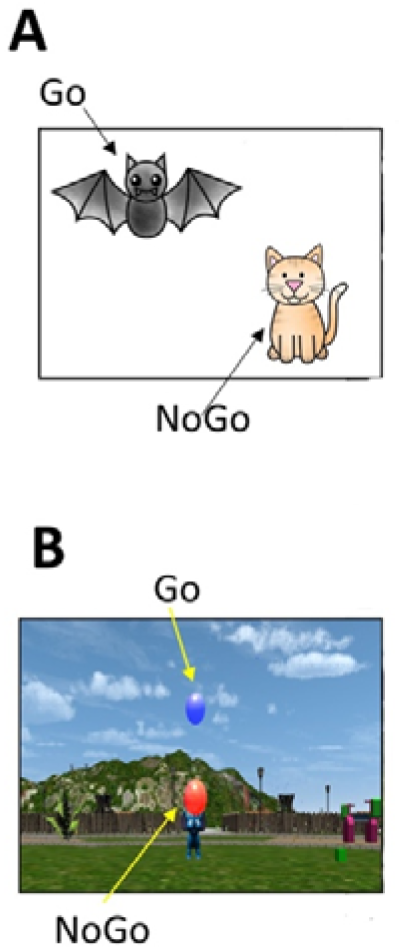
Experimental protocol. Children performed a computerized version of the Go/No-Go task sitting in front of a computer (A) and a VR version, while freely moving in a VR CAVE (B). In the CAVE children wore the fNIRS cap in conjunction with motion tracking markers and custom-made shutter glasses.

*VR Go/No-Go Task (VR).* The VR version of the task was performed in the Peltz VR CAVE at the ToddlerLab, Birkbeck, University of London (Figure 1 B). The CAVE is an immersive VR space made of a four-sided custom designed projection system (Mechdyne Corporation). The projection space includes a front wall (4.3 x 2m), two side walls (2.4 x 2m), and the floor (4.3 x 2m). Two blended single chip laser projectors overlapped by 65% (resolution of 2716 x 1528 pixels; total resolution= 3297x1528 pixels) display images onto the front and floor walls; a single laser projector (resolution 2716 x 1528 pixel) is used to present images on the side walls. Children wore an in house-built 3D printed model of child-sized pair of LCD shutter classes for active stereo viewing at 30 Hz that increases the immersive experience of participants in the virtual scene. Four six-degree-of-freedom optical motion tracking cameras (Vero 1.3 X, Vicon) at the corners of the CAVE tracked the orientation and movements of the children’s head and right hand. Head tracking was achieved by reflective markers mounted onto the shutter glasses and allowed for the virtual scene to be reoriented according to the participant’s position. Reflective markers were also attached onto a child-sized glove to track the right-hand movement which was used as a means of interaction with the virtual objects. The right hand was tracked for all participants regardless of handiness as the task did not require fine motor skills. The same glove was virtually presented in the virtual environment and the position was updated in real-time based on the participant’s hand position and movements.

In this version, participants found themselves in a virtual playground. An elephant-shaped bubble machine was located at the center of the playground and generated virtual bubbles of two different colors: blue (Go trials) and red (No-Go trials). Children were asked to pop the blue bubbles (Go) but not the red bubbles (No-Go trials) by means of the motion-tracked glove. The task started with practice trials to make the help the child acclimated to the virtual space and was comfortable and confident in how to use their right hand to pop the bubbles.

Both the VR and CB Go/No-Go tasks were block-designed, with 6 Go-only blocks and 6 Go/No-Go (Mixed; 50% Go trials, 50% NoGo trials) blocks. A total of 120 trials were split in 90 Go-only trials and 30 NoGo trials across the blocks. Each block had between 9 to 11 trials each; each trial was presented on screen or the CAVE for 2 seconds with an intertrial interval of maximum 1 second. Task blocks were spaced by rest periods with a randomized duration between 8 and 12 seconds; in the computer-based task children were asked to look at a fixation cross at the center of the screen. To replicate this in the CAVE, they were asked to fixate a star appearing on the elephant’s trunk. Each task took between 6 to 8 minutes to complete. Each task took between 6 to 8 minutes to complete. Go-only and Mixed blocks were alternated. The order of the CB and VR tasks was counterbalanced across participants.

### fNIRS data acquisition

Two wearable and wireless continuous wave fNIRS devices (Brite MKII, Artinis Medical Systems BV, Netherlands) were combined onto the same cap to measure the concentration changes of HbO_2_ and HHb while children carried out both the VR and the CB Go/NoGo tasks. Each instrument is equipped with 10 light sources, emitting light at 760 nm and 840 nm, and 8 detectors, sampling intensity data at 25 Hz. Optodes were arranged in the configuration shown in Figure 2A, providing 44 long separation channels (LSC) and 4 short separation channels (SSC).

**Figure 2.**
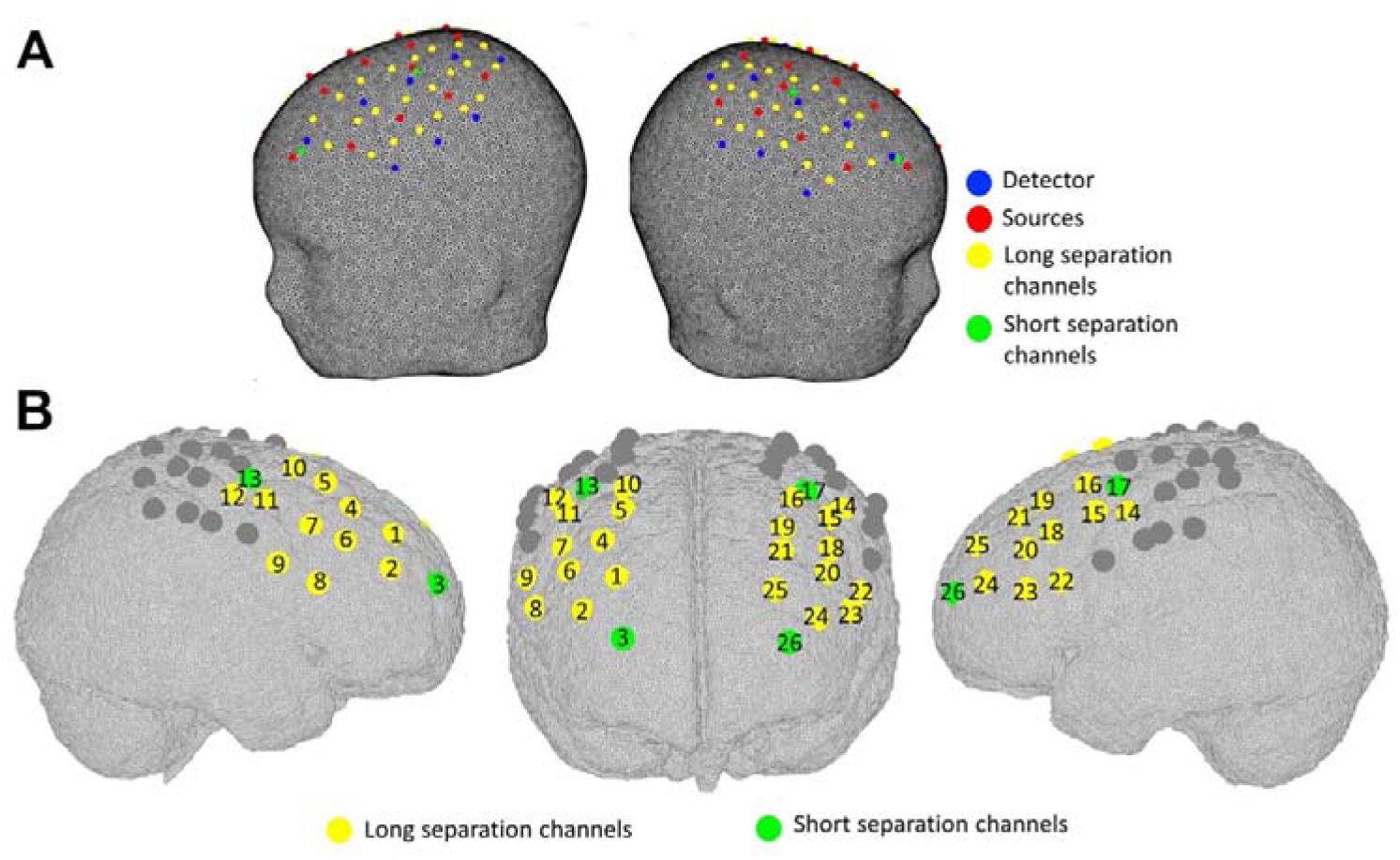
fNIRS probe configuration. Optodes’s arrangement (A) and corresponding channel numbers (B). Short and long separation channels are marked in green and yellow respectively. Channels excluded from the analysis are shown in grey.

The source-detector separation for the LSCs was set at 2.5 cm and 1 cm for the SSCs. Sources and detectors covered the dorsolateral prefrontal cortex (dlPFC) and the motor cortex bilaterally; given that the focus of this work is on the neural correlates of inhibitory control, we focused our analyses on the 26 channels covering the right and left dlPFC (Figure 2B).

To achieve a reliable placement across participants, the cap was aligned to the Fp1 and Fp2 landmarks of the 10-20 electrode placement system. A short 10s long frontal video of the participant was recorded to perform the co-registration of the fNIRS array onto a common template (see the fNIRS data pre-processing and analysis section below).

### fNIRS data pre-processing and analysis

For each participants, the locations of the optodes and channels were co-registered onto a common 5-year-old MRI template from the Neurodevelopmental MRI Database of the University of South Carolina (http://jerlab.psych.sc.edu/NeurodevelopmentalMRIDatabase/) using the procedure described in Bulgarelli et al. (2023). Briefly, we have 3D printed the MRI template to create our head model and to specify the ideal placement of the cap; the head model coordinates of sources and detectors as well as anatomical landmarks (Nasion, Inion, Cz, right and left pre-auricular points, Fp1, Fp2, Fpz, F7, F8, O1, O2) were recorded using a 3D magnetic digitizer (Fastrak, Polhemus). These were used as an input to STORM-Net (https://github.com/yoterel/STORM-Net) for the off-line stage of the algorithm. The frontal videos of the participants were then used for the online step to estimate the position of the child’s optodes and landmarks based on the displacement respect to the ideal head model (Erel et al., 2020). The MRI template was segmented into 5 tissues (scalp, skull, cerebrospinal fluid, grey matter (GM), white matter) using FSL and a volumetric multilayer mask was then created using routines from the DOT-HUB toolbox (https://github.com/DOT-HUB). A tetrahedral volumetric mesh and a GM surface mesh were generated; the optodes’ coordinates for each participant were co-registered onto the scalp mesh first via affine transformation and then projected onto the GM surface mesh. The same procedure was applied to the head model coordinates. The LPBA40 atlas was used on the head model data to identify the anatomical locations of all channels; the anatomical labels of the channels and their percentage of overlap are reported in Table 1.

**Table 1.**
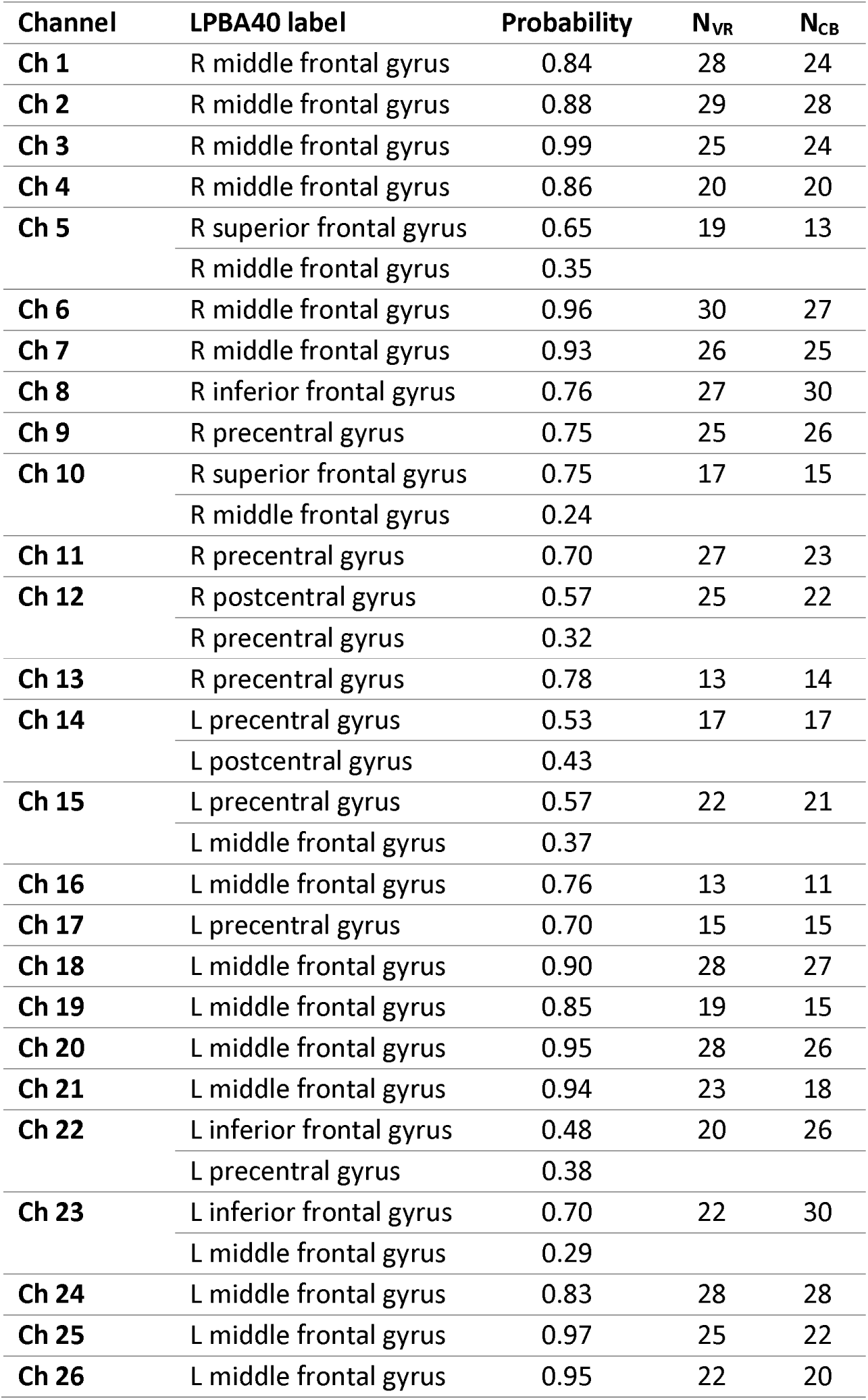
Anatomical locations of the fNIRS channels. The anatomical areas (LPBA40 atlas) and the corresponding atlas-based probabilities for each channel are included. Only probabilities greater than 20% are listed. Number of participants contributing to each channel for the VR (N_VR_) and CB (N_CB_) tasks because of cap placement and data quality are also listed.

Raw intensity fNIRS data were first visually inspected to identify noisy channels to further exclude from following analyses; channels with no clear heart rate peak or detector saturation or severely impacted by motion artifacts were excluded. Table 1 includes the number of participants that contributed with good quality data for each channel. Intensity signals were then pre-processed using Homer2 (Huppert et al., 2009). More precisely, two separate pipelines were applied to both the VR and CB tasks (Figure 3) in order to compare the impact of SSR.

**Figure 3.**
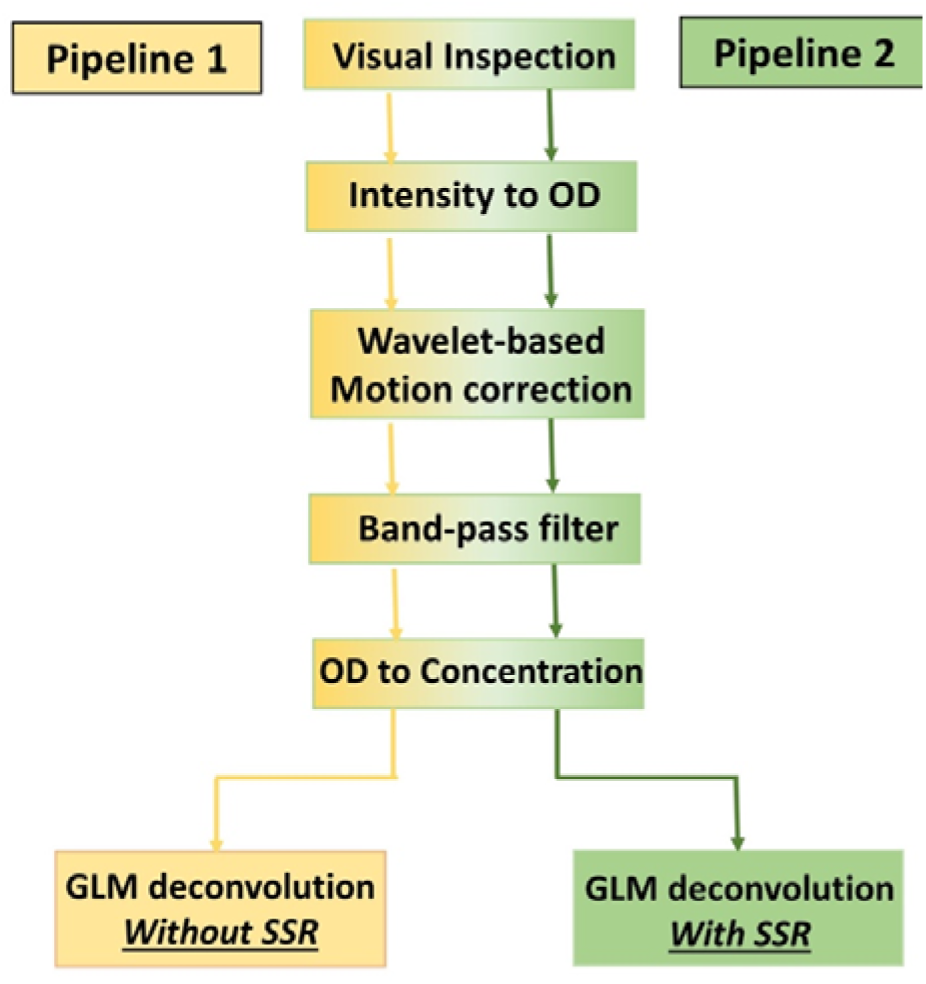
Analysis pipelines.

In both pipelines, raw fNIRS data were first converted into changes in optical density (*hmrIntensity2OD*) and corrected for motion artifacts using the wavelet-based algorithm (iqr=0.8, *hmrMotionCorrectWavelet* (Molavi & Dumont, 2012)). Optical density signals were band-pass filtered (Fc=[0.01 0.1] Hz; *hmrBandpassFilt*) and converted into changes in HbO_2_ and HHb through the modified Beer-Lambert law (DPF= 5.5, 4.7; *hmrOD2Conc* (Scholkmann & Wolf, 2013)).

Participants were included in the analysis if they had at least 50% of the fNIRS channels of good quality and at least 3 blocks with performance >50% for both the Go-only and Mixed blocks. A General Linear Model-based deconvolution approach was then used to estimate the hemodynamic responses to the Go- and Mixed blocks, for each participant, channel and chromophore. The hemodynamic responses were extracted using a set of gaussian basis functions with standard deviation and temporal spacing of 1.5 s in the time window [-2 28] s around each onset of the included blocks; for pipeline 2, the design matrix included the short separation channel with the highest correlation to each long separation channel (Zhao et al., 2020); no SSC regression was performed in pipeline 1 (trange = [-2 28], glmSolveMethod = 1, idxBasis = 1, paramsBasis = [1.5 1.5], rhoSD_ssThresh = 0 for pipeline 1 and 1.5 cm for pipeline 2, flagSSmethod = 1, driftOrder = 0, flagMotionCorrect = 0; *hmrDeconvHRF_DriftSS*).

For both the VR and CB tasks, the area under the curve (AUC) in the estimated responses to the Go and Mixed blocks were computed within a time window from 15 to 25s post task onset; this was done for each pipeline and for both HbO_2_ and HHb of each participant and used for the group level statistics. Channel-wise one-sample t-tests were run on the group AUCs for each pipeline to test whether there is a statistically significant (p<0.05) larger increase in HbO_2_ and a larger decrease in HbR in the Mixed blocks versus the Go blocks. One-sample t-tests were also used to test whether there were significant hemodynamic changes in the SSCs in response to either the Mixed or the Go blocks. Results were corrected for multiple comparisons using the False Discovery Rate method (FDR; (Benjamini & Hochberg, 1995)).

## Results

### Optimization of the preprocessing pipeline

Initial visual inspection of the raw intensity data revealed that some children exhibited signal changes with a periodicity of 10s corresponding to Mayer waves, similarly to what it has been reported in adults (Yücel et al., 2016). These were clearly visible from the raw intensity time series, both in the long separation channels and in the short separation channels. In Figure 4, we show an example of raw signals at the two wavelengths for a long (channel 12) and a short separation channel (channel 13) for one child. Besides the heartbeat, the Mayer wave component can be clearly distinguished for the VR task (Figure 4 A) and, to a lesser extent, for the CB task (Figure 4 B), suggesting that Mayer waves may be present both when fNIRS data are recorded from children who are standing and when they are sitting.

**Figure 4.**
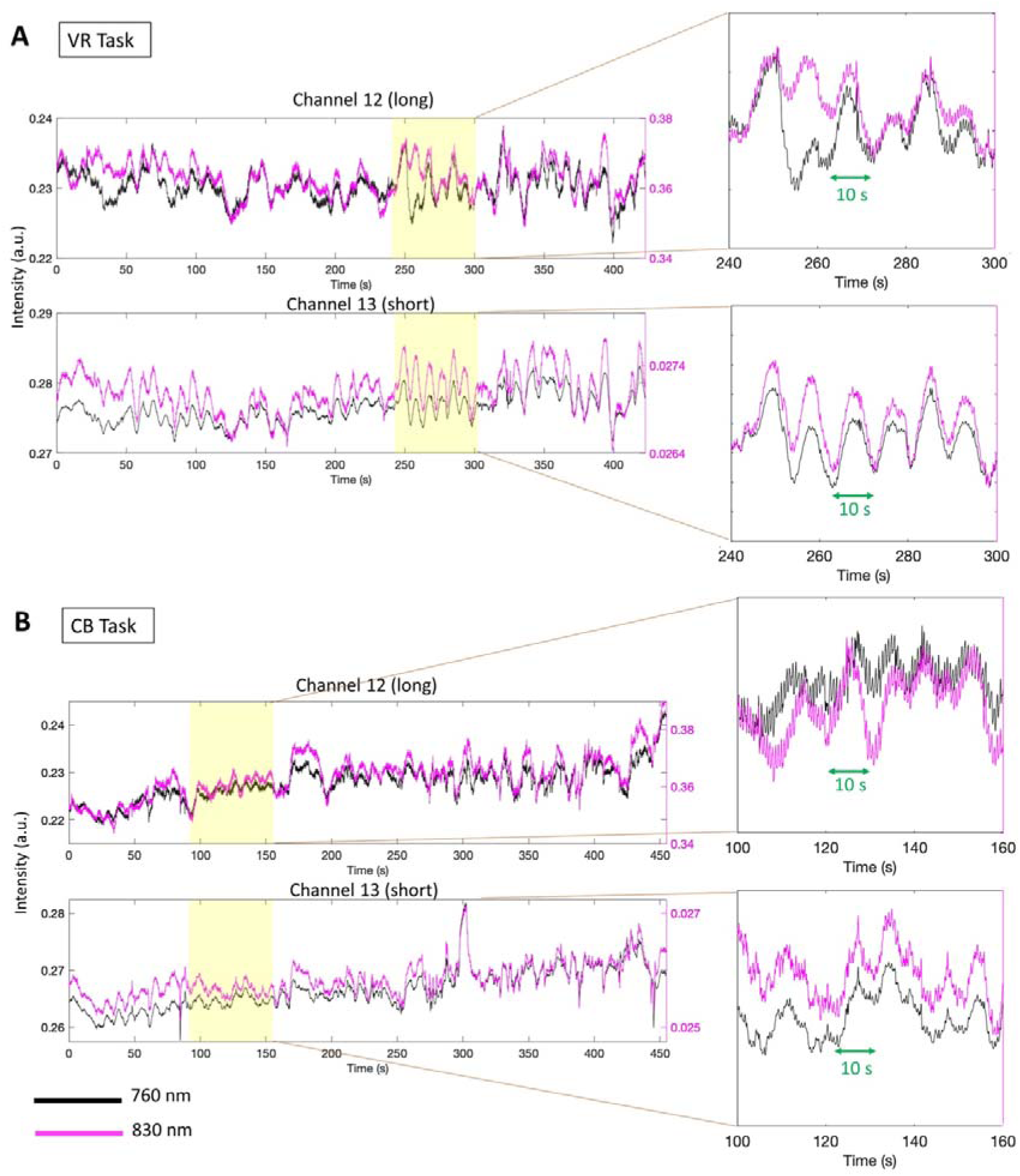
Examples of raw fNIRS data. Panel A and B show the raw intensity signals at the two wavelenghts for a long and a short separation channel of one participant, for the VR task (A) and the CB task (B). Periodic 10 s components corresponding to Mayer waves can be observed in all channels.

When using common low pass cut-off frequencies of 0.5 Hz (Yücel et al., 2021) in conjunction with a high pass filter at 0.01 Hz in pipeline 1, the 10 s modulations related to the Mayer waves are still visible in the block-averaged responses, especially for HbO_2_. Figure 5 (dotted lines) shows the resulting hemodynamic responses averaged across the VR Go blocks for channel 12 and channel 13 presented in Figure 4 A. It is important to notice that similar haemodynamic changes occur both in the long and in the short channel, with amplitude changes of approx. 0.5 μMol/L for HbO_2_.

**Figure 5.**
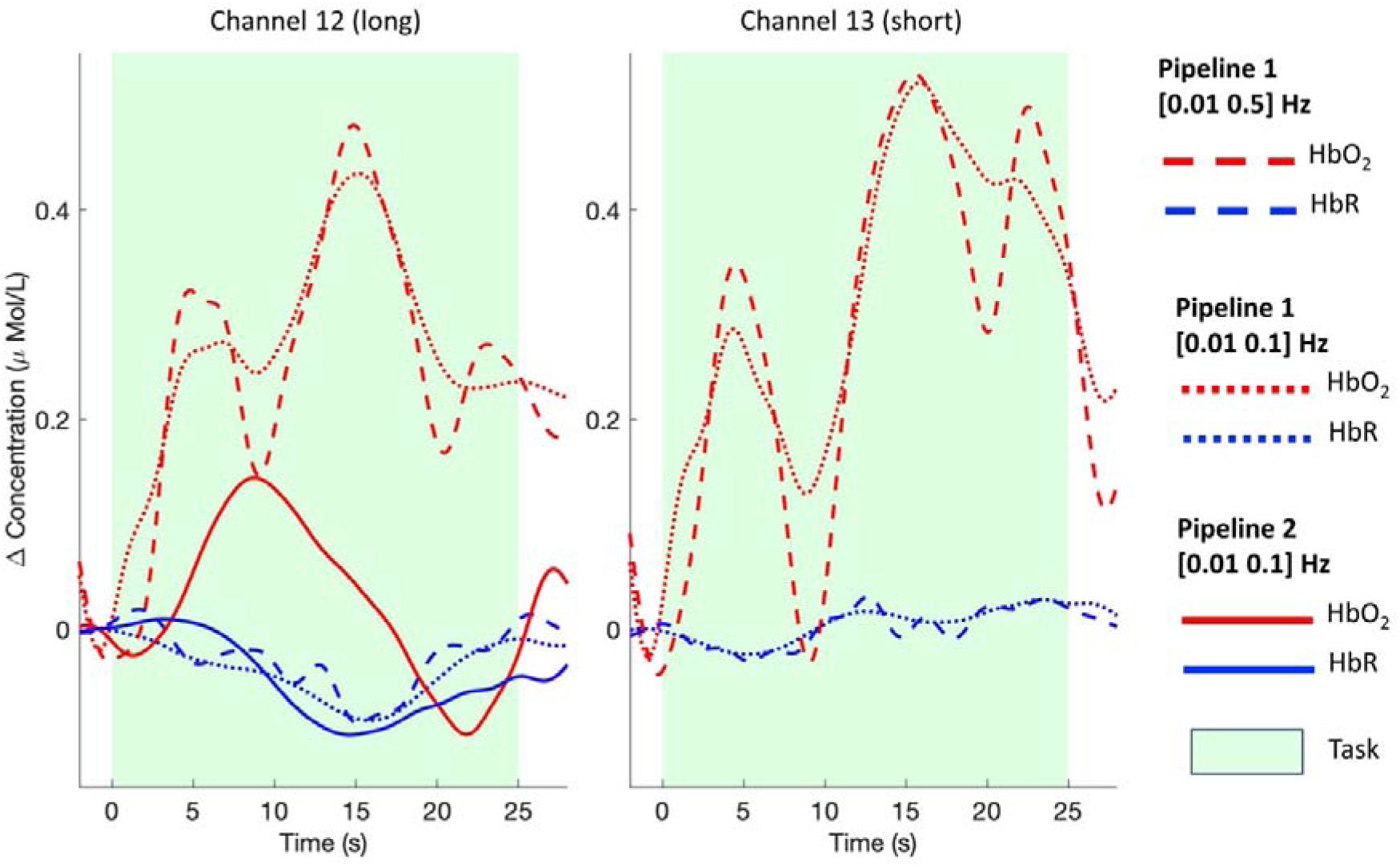
Impact of different pipelines on the reduction of physiological components. The estimated task-evoked hemodynamic responses with different pipelines are presented for the Go-only condition for one long and short separation channel from one participant. The use of a band pass filter only, either with a wider (dashed lines) or narrower (dotted lines) passband are not as effective as the combination of SSR with a narrower band pass filter (solid lines) in reducing the Mayer waves modulations in the signals.

However, when reducing the pass-band of the filter to [0.01 0.1] Hz, such modulations are attenuated (Figure 5, dashed lines). Therefore, even though Mayer waves cannot be simply removed by low pass filters (Yücel et al., 2016), a low pass cut-off frequency closer to the Mayer waves fundamental frequency (0.1 Hz) can help in attenuating the effect of such component. It is worth mentioning that these parameters are appropriate for our task design with a task frequency of approx. 0.03 Hz, with the [0.01 0.1] Hz including 3 harmonics of the task frequency. In case of faster task designs, careful considerations should be made in narrowing the pass-band of the filter (Pinti et al., 2019). When performing short channel regression following the [0.01 0.1] Hz filter in pipeline 2, the Mayer waves modulations are significantly reduced (Figure 5, solid line), particularly in HbO_2_, suggesting that for our type of experimental context and population the combination of these preprocessing steps seems to be optimal in reducing the impact of physiological noise.

### Extracerebral changes and impact of SSR

At the group level, significant changes were found in the short separation channels (Figure 6). For the VR task (Figure 6 A), we found a significant decrease in HbO_2_ in Channel 3 (*t_ch3_*=-2.07; p<0.05 uncorrected) and a significant increase in HbR in Channel 13 (*t_ch13_*=2.38; p<0.05 uncorrected) for the Go-only condition, and a decrease in HbO_2_ in Channel 26 (*t_ch26_*=-2-26; p<0.05 uncorrected) for the Mixed condition. For the CB task (Figure 6 B), a significant decrease in HbO_2_ in Channel 3 (*t_ch3_*=-2.07; p<0.05 uncorrected) and a significant increase in HbR in Channel 13 (*t_ch13_*=2.38; p<0.05 uncorrected)

**Figure 6.**
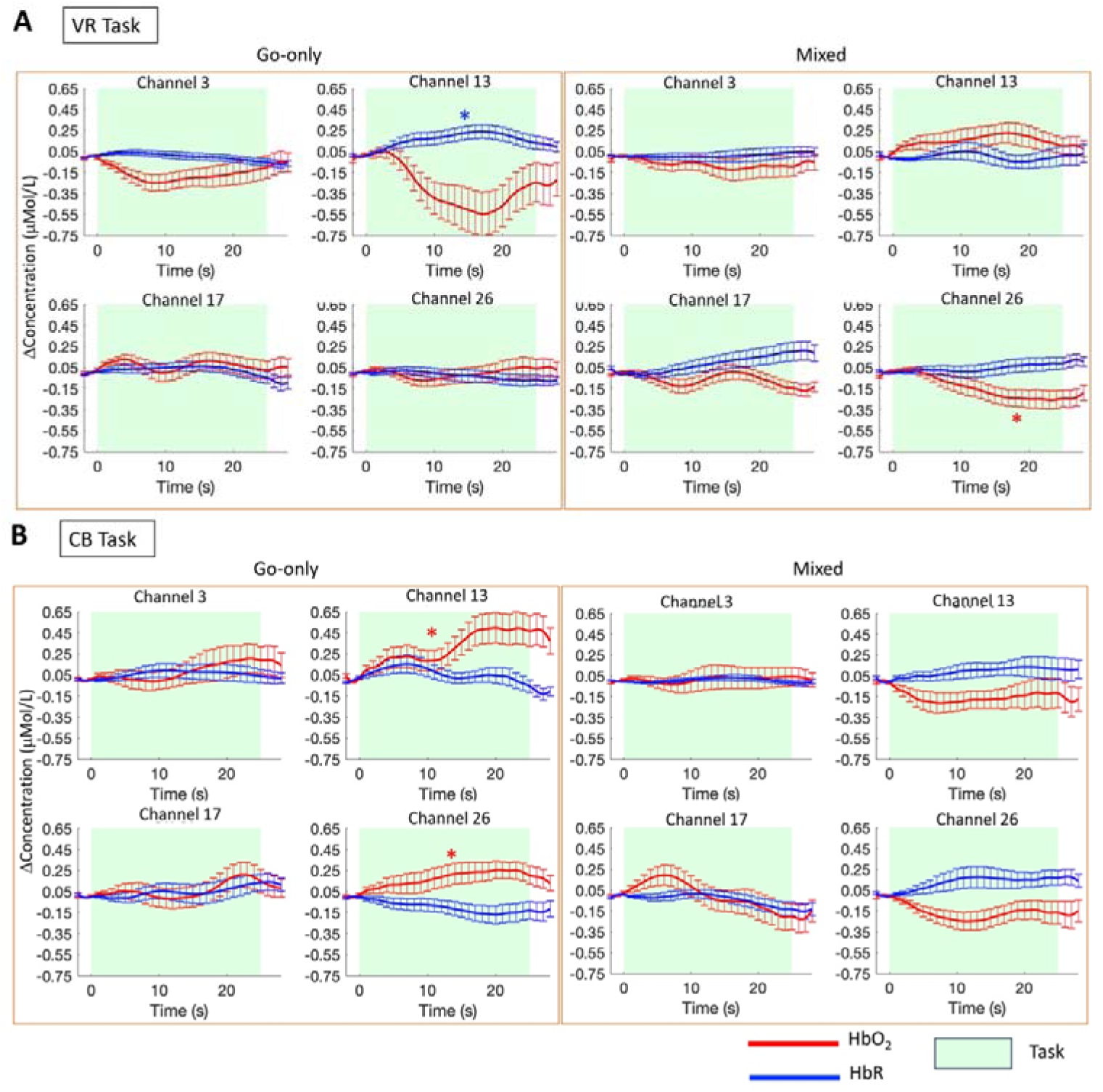
Group averaged (mean ± std.err.) HbO_2_ (red) and HbR (blue) responses in the short separation channels. Task-evoked haemodynamic responses are show for the both the Go-only and Mixed conditions and for the VR (A) and CB (B) tasks.

To test the effect of the regression of short separation channels on the group level statistics, we ran channel-wise one-sample t-tests on the group AUCs for our contrast of interest (Mixed > Go-only) for HbO_2_ and HbR for pipeline 1 and pipeline 2. This was done for both the VR and CB task. Group-level *t*-maps are shown in Figure 7 and Figure 8 for the VR and CB experiments respectively.

**Figure 7.**
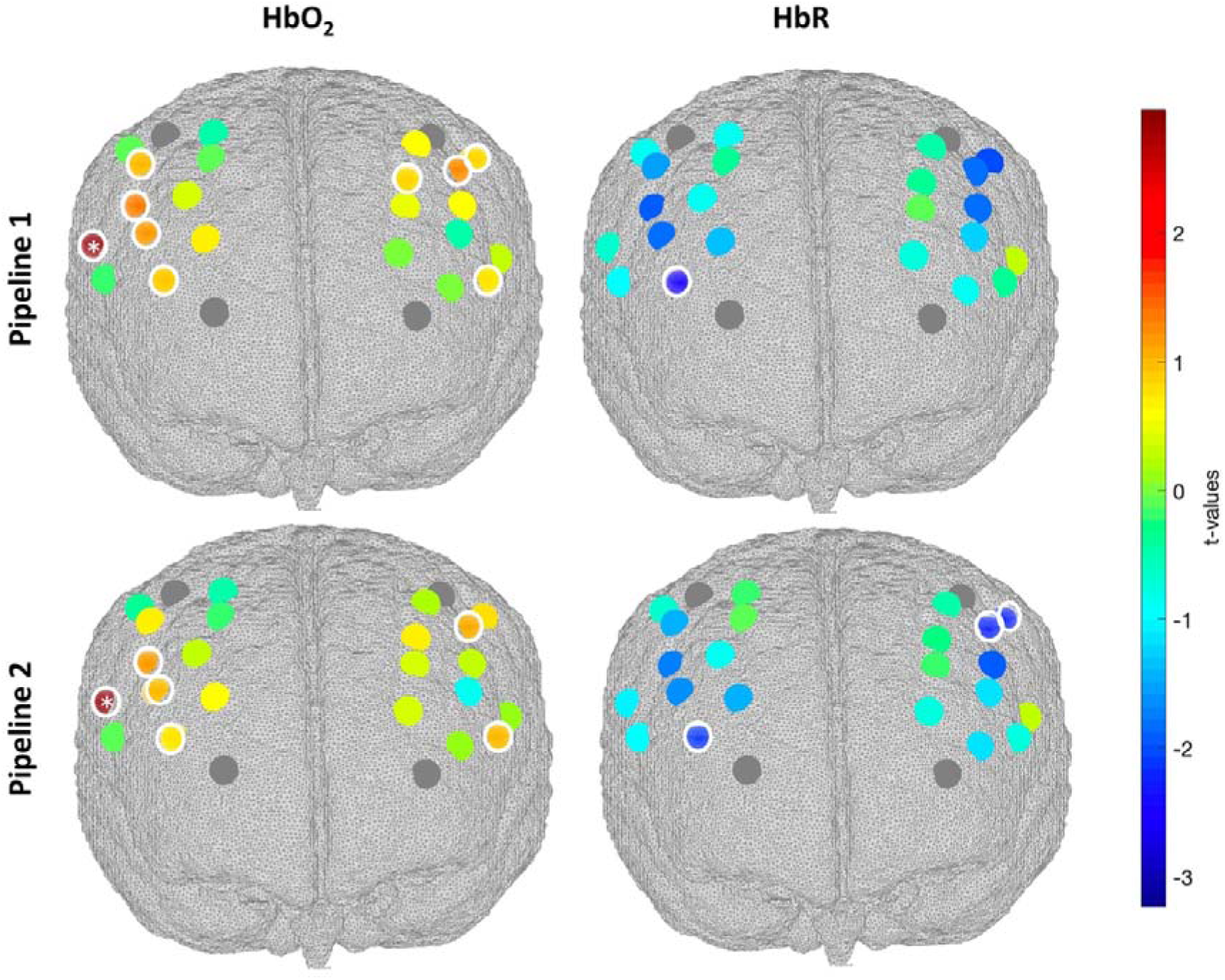
Group level t-values maps for the contrast Mixed>Go-only for the VR task. Statistically significant channels at p<0.05 are circled in white. Channels surviving FDR correction are marked with asterisks. A positive t-value corresponds to a HbO_2_ increase and a HbR decrease; a negative t-value corresponds to a HbO_2_ decrease and a HbR increase.

**Figure 8.**
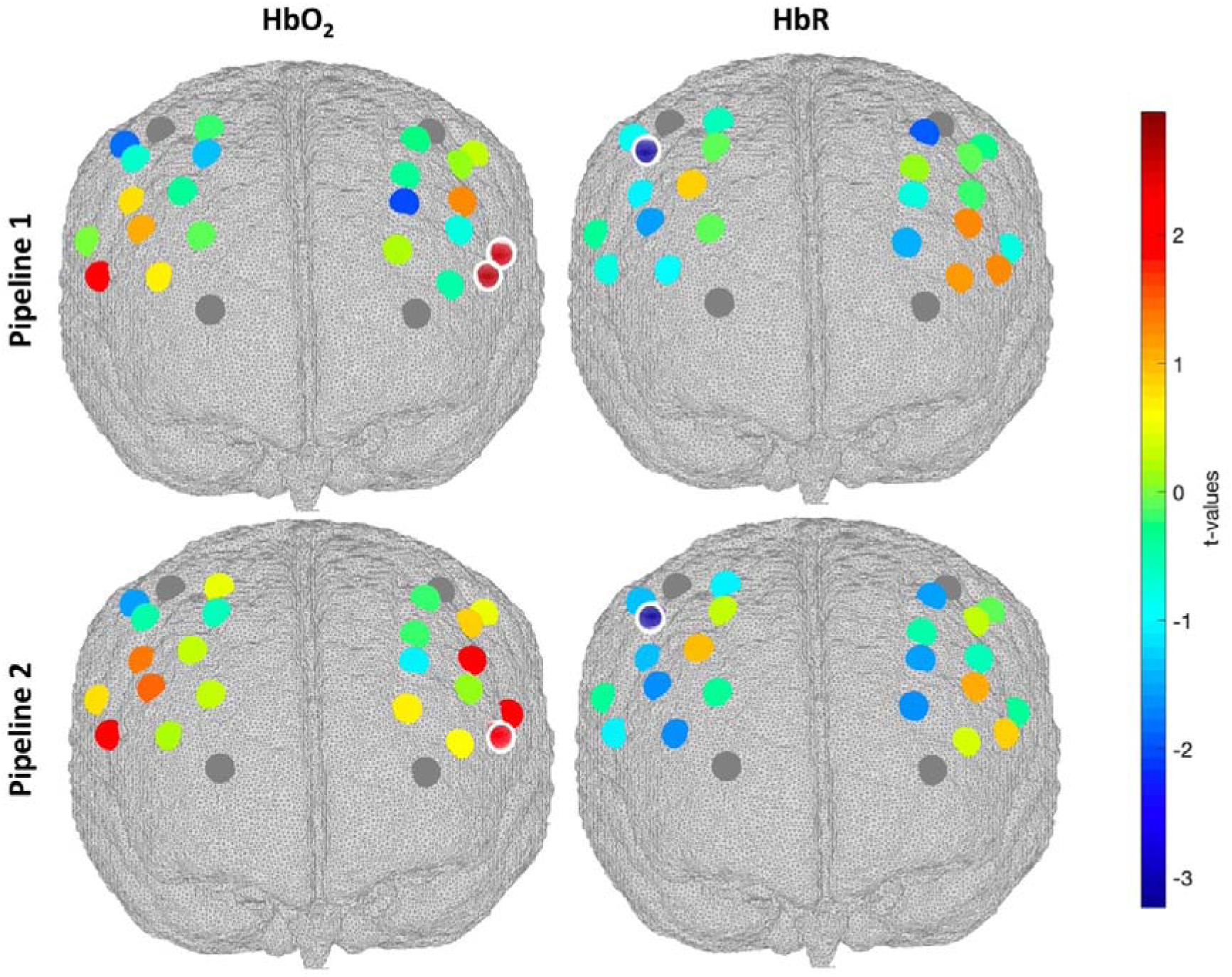
Group level t-values maps for the contrast Mixed>Go-only for the CB task. Statistically significant channels at p<0.05 are circled in white. Channels surviving FDR correction are marked with asterisks. A positive t-value corresponds to a HbO_2_ increase and a HbR decrease; a negative t-value corresponds to a HbO_2_ decrease and a HbR increase.

For the VR task, significant changes (p<0.05) in HbO_2_ and HbR can be observed in the channels covering the middle frontal, precentral and inferior frontal gyri, although only Channel 9 (inferior frontal gyrus) survives the FDR correction for HbO_2_. Looking at the general and uncorrected patterns of brain activity, more widespread activations are found when using pipeline 1, with nine significant channels (p<0.05) for HbO_2_. Pipeline 2 yields more localized activations, with six significant channels for HbO2 (p<0.05). For HbR, the regression of short channels in pipeline 2 increases the statistical significance of two more channels, further maximizing the overlap between HbO_2_ and HbR that better reflects the expected dynamic of brain activity, i.e. increase in HbO_2_ and decrease in HbR.

Similarly for the CB task, we found significant changes (p<0.05) in HbO_2_ and HbR in the middle frontal, precentral and inferior frontal gyri, although no channels survive the FDR correction. Similarly to the VR case, the regression of short channels results in more localized changes in HbO_2_, but no effect is observed for HbR. This suggests that physiological changes might affect brain haemodynamics more on standing (VR case) versus seated (CB case) participants, and that in case of computerized task HbR is more robust to systemic interferences as previously reported (Tachtsidis & Scholkmann, 2016).

In general, in both tasks, the superficial signal regression led to a reduction in the amplitude of HbO_2_ and increase of that HbR. An example of the resulting group responses to the contrast Mixed-Go-only is shown in Figure 9 for two representative channels, for both the VR and CB tasks.

**Figure 9.**
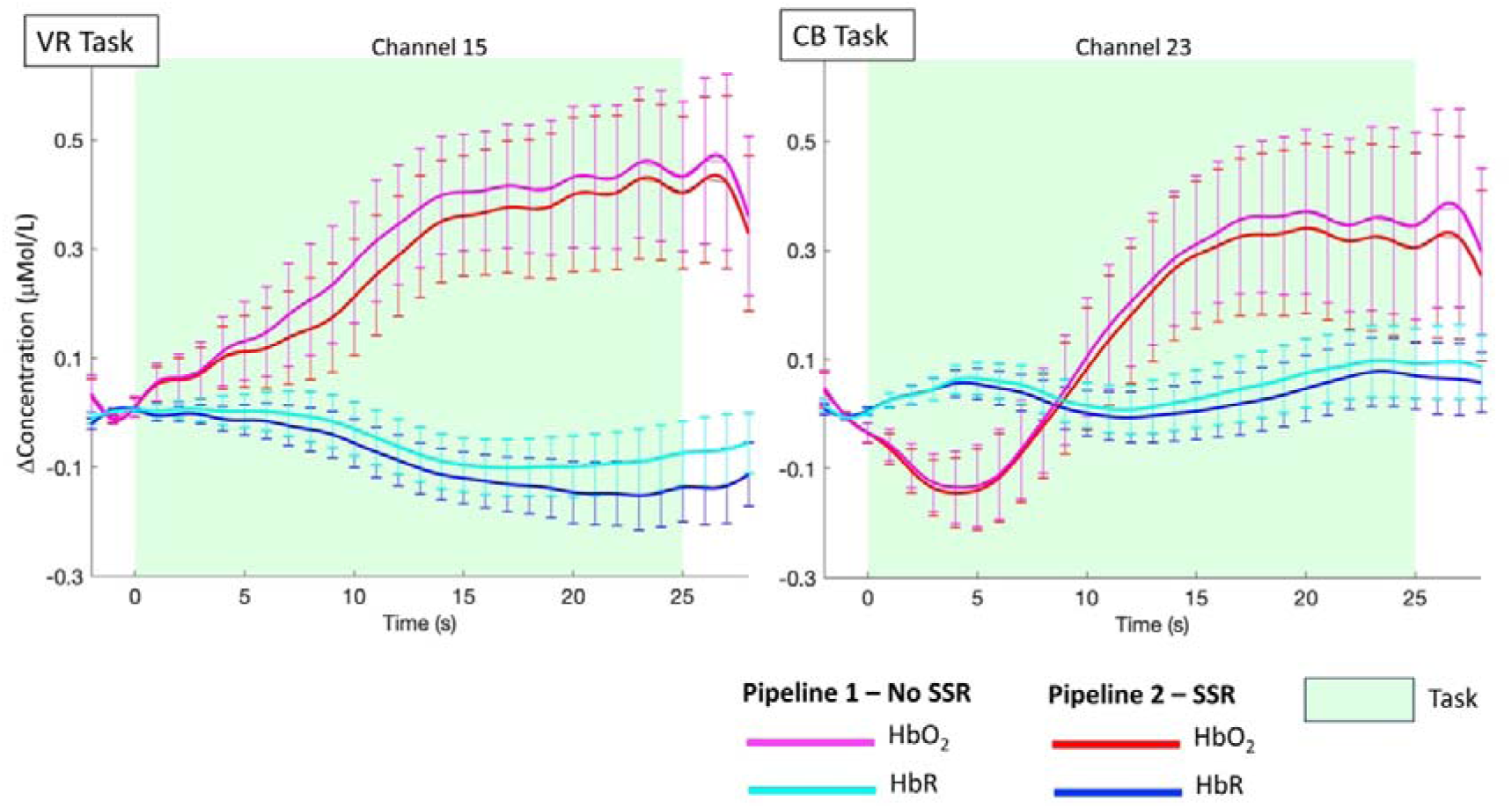
Single subject task-evoked average responses (mean ± std.err.) for the contrast Mixed−Go-only. The resulting HbO_2_ and HbR responses from two representative channels of one participant are shown both without (magenta and cyan lines) and with SSR (red and blue lines) for the VR and CB tasks. SSR leads to a decrease in HbO2 amplitude and an increase in HbR changes.

## Discussion

In this work, we investigated for the first time in toddlers and preschoolers if significant haemodynamic and oxygenation changes occur in the extracerebral layers of the head and whether the removal of such components has an impact on the group level statistics when localizing haemodynamic responses in the cerebral compartment. Specifically, we explored whether the superficial contamination on the cortical brain fNIRS signals was stronger when data are recorded from a developmental sample during a cognitive task while freely moving compared to sitting. To this goal, we have collected fNIRS data from 3-to-7 year-olds during a Go/No-Go task performed while freely moving in an immersive VR CAVE space (VR task) and while sitting in front of a computer (CB task). We have applied two different analysis pipelines (Figure 3) to test the effect of superficial signal regression on the group level statistics.

Our initial investigations suggested that similar physiological components as adults’ fNIRS data can be observed in children fNIRS data as shown by visible Mayer waves in the raw intensity data (Figure 4) alongside other known components such as heart rate, slow trends, and motion artifacts. If these are not accounted for, they can propagate along the analysis pipeline, possibly ending up affecting the outcome of the statistical analysis. For example, in Figure 5 we showed that if appropriate approaches, i.e., band-pass filtering along with SSR, are not used, Mayer waves components are still clearly visible in the recovered haemodynamic responses, leading to an inflation of the amplitude changes of HbO_2_ and hence to false positives (Tachtsidis & Scholkmann, 2016). Similar to previous studies in adults (Gagnon et al., 2012; Noah et al., 2021; Yücel et al., 2016), we found significant changes in the short separation channels (p<0.05 uncorrected; Figure 6) both in the VR task and in the CB task, indicating that physiological contamination can occur not only when children are standing or moving around, but also when they are performing standard computer-based tasks. Such systemic changes also seem to be heterogenous across the scalp as previously reported in adults (Gagnon et al., 2012; Wyser et al., 2020). Hence, the use of multiple short separation channels can be highly beneficial in this population as much as it is in adults.

Our exploratory analyses confirmed that the Go/No-Go task elicited significant brain activity in the expected regions of interest within dlPFC (Munakata et al., 2011; Wu et al., 2023) and revealed that the regression of short separation channels has an effect on the group level statistical maps both for both the VR and CB tasks (Figure 7 and 8). In particular, pipeline 2 (with SSR) led to more localized activation patterns compared to pipeline 1 (without SSR) for HbO_2_. The improvement in the spatial localization has also been reported by previous fNIRS studies on adults (e.g., (Noah et al., 2021; Yücel et al., 2015)). SSR also increased the statistical significance of HbR (Figure 7), further improving the detection of brain activity (i.e., increase in HbO_2_ and decrease in HbR). The increase in spatial localization per effect of SSR was more pronounced for the VR task (Figure 7) than for the CB task (Figure 8), suggesting that, in agreement with our hypothesis, systemic interferences might have a larger impact when fNIRS data are recorded in more physically active conditions (Pinti et al., 2015, 2018). Nevetheless, SSR influences group level results also for the CB experiment, even though to a lesser extent (Figure 8). Therefore, we also recommend the use of short separation channels in instances where fNIRS data are recorded from children in typical non-mobile neuroimaging contexts. Across both tasks, most of the changes post SSR can be observed in the HbO_2_ signal, in agreement with previous work demonstrating that oxyhemoglobin is more sensitive to physiological interferences than HbR (Kirilina et al., 2012).

In summary, our preliminary work represents the first report of the occurrence and the influence of scalp blood flow changes on the estimation of brain activity through fNIRS in toddlers and preschoolers. As such, it provides initial first evidence that: (1) significant changes in scalp haemodynamics can occur in toddlers and preschoolers; (2) extracerebral changes seems to be stronger when children perform cognitive tasks while moving versus when they are sitting; (3) SSR can influence the outcome of group level statistics by improving spatial localization of hemodynamic responses; (4) short separation channels can be highly beneficial in 3 to 7 year-olds both in mobile and non-mobile experimental setups. However, limitations to this work should be considered. First, the sample size was not always ideal for all the channels (Table 1). This depended on whether the optodes were probing brain activity on hairy regions, that worsened the optical coupling in some participants; when collecting data on young children, coupling optimization procedures cannot always take place or need to be done very quickly to avoid kids becoming uncompliant. In addition, the VR and the CB tasks involved different body movements and changes in optical coupling from one task to the other, and re-optimization procedures could not always take place; for example, some children moved their head more during the CB task than the VR task. Collecting data from a larger cohort of children could help in mitigating these issues in future studies. In addition, we have not used additional monitors of systemic physiology (e.g., ECG, respiration, blood pressure) that could have helped to better assess the presence of significant task-related physiological changes and further improve the robustness of fNIRS signals. This was partly related to practical challenges in using several pieces of equipment on young children as in our study they were already wearing the fNIRS cap, a motion tracking glove, and head mounted shutter glasses/head tracking (Figure 1). Nevertheless, previous work has shown that a Systemic Physiology Augmented fNIRS (SPA-fNIRS) approach (Yücel et al., 2021; Zohdi et al., 2021) can be highly beneficial in denoising the fNIRS signals from systemic interferences. This would be worth exploring in children in future studies. Alongside physiological monitors, in our future work we plan on using signals coming from motion tracking, such as head movements, as additional nuisance regressors. In fact, von Lühmann and colleagues (2020) (Von Lühmann et al., 2020) showed that combining short separation channels with accelerometer signals on the head can significantly improve the recovery of the hemodynamic responses. Finally, we chose a source-detector separation of 1 cm due to the physical constraints of the optodes size; even though young children seem to have similar sensitivity profile to adults (Fu & Richards, 2022) and similar head sizes, it is possible that part of the signal of the short separation channels is of cortical origin; therefore, shorter source-detector separations might be better suited for SSR in children (Emberson et al., 2016) if hardware allows.

### Conclusion and methodological recommendations

In this study we have provided first evidence that superficial signal contamination of fNIRS data also occurs in younger children. Interestingly, our results indicate that children from 3-to-7 year-olds are more similar to adult populations rather than infants in terms of non-neural components in the fNIRS signals. In fact, previous work from Emberson and colleagues (2016) showed that the removal of superficial vasculature signals did not have a significant impact on group level statistics in 6-month-old infants. Although, it is unclear whether other, more active or physiologically arousing paradigms might show greater contribution of superficial vasculature signals even in infants. The assumption that the fNIRS signal is a mixture of components of both neuronal and non-neuronal origin (Tachtsidis & Scholkmann, 2016) seems to be valid in children as well and can potentially lead to false positives and false negatives at the statistical inference stage. Therefore, we suggest that removing the contribution of superficial signals to the long fNIRS channels is an important step in the analysis pipeline of children’s data, especially when fNIRS signals are recorded in experimental contexts that elicit stronger physiological changes (e.g., standing, walking, full body movements). Future work will be important to further explore how other cognitive tasks and experimental settings modulate the impact of systemic interferences in children as well as the feasibility of SPA-fNIRS in these populations.

Task design, array co-registration, and the preprocessing pipeline were optimized for children to maximize the robustness of the results. Based on our results, below we provide preliminary recommendations as a first step to advance the field of children fNIRS neuroimaging. Figure 10 shows some of the steps that we advise to follow to enhance the robustness of the results of fNIRS data recorded on children and particularly in ecological settings.

**Figure 10.**
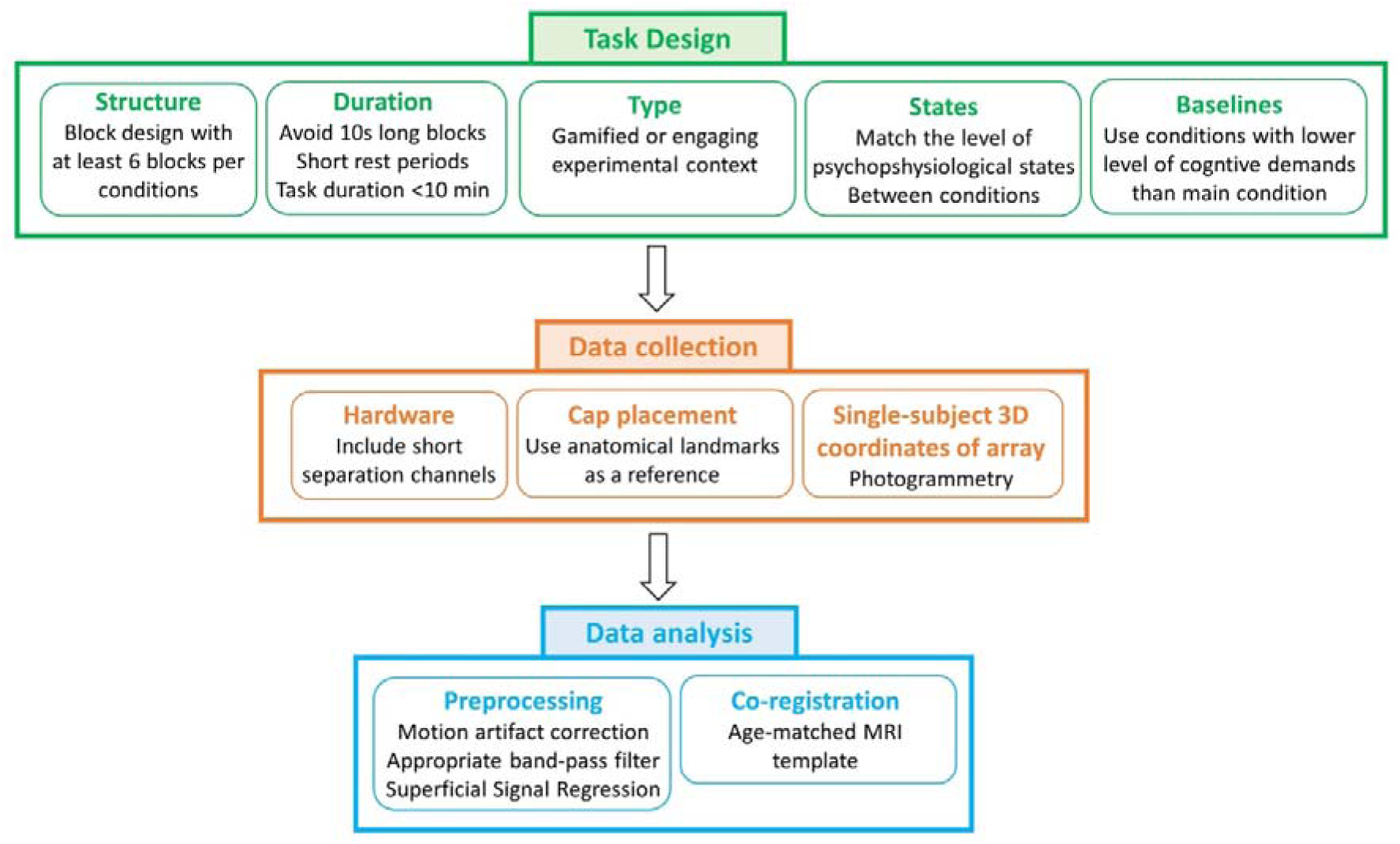
Summary of methodological consideration for robust children neuroimaging experiments.

*Methodological considerations for task design.* Tasks were designed to meet different requirements.

First, it is well known that children neuroimaging can present several challenges due to their compliance to neuroimaging experimental requirements, particularly in preschoolers (Barkovich et al., 2019). The CAVE represents a much more engaging setting for children, increasing their compliance together with fNIRS being a child-friendly technique (Turk-Browne & Aslin, 2024). Hence, longer tasks with a larger number of trials that increase the statistical power of the study may be potentially suitable. However, in this work a conventional computerized task was needed for the validation of the novel VR naturalistic task and to evaluate the differences in physiological interferences between the two. To maximize children’s cooperation and participant’s retention to two experiments, both tasks were kept to a relatively limited duration of 6 to 8 minutes each while ensuring a good number of trials. In particular, 6 blocks per condition were used to ensure that in case of performance- or attention-based blocks exclusion, the chances of ending up with at least 3 blocks per condition were high (NOTE: a threshold of minimum of 3 blocks for participant’s inclusion is often used in fNIRS developmental research (Lloyd-Fox et al., 2017)).

Second, to maximize the signal-to-noise ratio of the data, we chose block-designed tasks rather than an event-related design as the latter requires a larger number of events and usually elicits lower contrast haemodynamic responses above to the noise level (Henson, 2015). Along these lines, block designs typically include repeated rest periods to allow the slow haemodynamic response to go back to baseline; however, it has been suggested that it is not fully appropriate to contrast the task to the rest blocks as it is unknown what the participant is really doing during that rest time and hence it might not be a realistic baseline. It is instead recommended to use low-level control conditions for experimental contrasts (Henson, 2015; Pinti et al., 2023). Therefore, our task was designed such that Mixed blocks were contrasted to the lower-level Go-only blocks. Short rest periods were added in between to avoid the overlap of the various haemodynamic responses but were not used for comparing conditions.

Finally, to help minimizing the impact of physiological interferences especially in the VR task, we avoided 10s-long blocks (Tachtsidis & Scholkmann, 2016) and the conditions (Go-only and Mixed) were planned in a way to have the same level of physical activity. In this way, when contrasting the Mixed blocks versus Go-only, some of the confounding effects may cancel out (Pinti et al., 2023). In the VR task, both conditions required the participants to stand and to move the arm in the same way; in the CB task, a button press was always needed.

*Methodological considerations for data collection.* To take into account task-evoked extracerebral changes, we suggest adding short separation channels to the fNIRS array, even if the experiment is performed while sitting. We recommend including at least one SSC although it has been previously demonstrated that scalp interference is not homogenous and, based on hardware availability, a larger number of SSC scattered around the head is advised (Gagnon et al., 2014).

In order to account for between-subjects variability in the location of the channels, the co-registration of the fNIRS array onto a common brain template is typically recommended, besides ensuring a reliable cap placement using anatomical landmarks as a reference. 3D magnetic digitizers are generally used in fNIRS (and EEG) studies to gather the 3D coordinates of the optodes and landmarks. However, these require the participant to stay still for several minutes and are also susceptible to electromagnetic interference from the environment (Pinti et al., 2023). Therefore, it may not be feasible to have children stay still for long periods of time. To overcome this barrier, here we used a photogrammetry-based method (STORM-net (Erel et al., 2020)e, but others are available (Hu et al., 2020; Jaffe-Dax et al., 2020; Mazzonetto et al., 2022)) that is compatible with children’s movements and does not require extra hardware except a smartphone camera.

*Methodological considerations for data analysis.* In terms of the preprocessing of fNIRS data, as demonstrated by previous studies, we recommend including motion artifact correction rather than trial rejection in the preprocessing pipeline (Brigadoi et al., 2014); this ensures that a larger number of trials can be retained which is particularly important when working with developmental populations where the number of good quality blocks might be limited. Following the procedure described in (Pinti et al., 2019), we recommend choosing a filter that is appropriate for the timings of the task and, when possible, use a narrower low pass filter with a cut-off frequency close to 0.1 Hz (the Mayer waves fundamental frequency) to help attenuating its impact. Here for example, we used a Butterworth band-pass filter within the range [0.01 0.1] Hz. To improve the localization of functional brain activity by minimizing the effect of systemic interferences, the regression of short separation channels from the long separation channels seems to be an important step also in young children and hence we advise to incorporate it within the analysis pipeline. Finally, to further improve the robustness of the inferences about the task-evoked haemodynamic activity, it is important to know the underlying anatomical location of the fNIRS channel. This allows the grouping of anatomically and functionally homogenous regions in the group-level analysis and can be achieved by co-registering the 3D coordinates of the optodes/channels onto a common brain template. Whilst co-registering the channels onto the subject-specific structural MRI can better capture the individual variation in cortical anatomy (Turk-Browne & Aslin, 2024), these are rarely available, especially for children; therefore, we recommend using an averaged age-matched templates that can be found available on open access repositories. For example, here we have used the Neurodevelopmental MRI Database of the University of South Carolina (http://jerlab.psych.sc.edu/NeurodevelopmentalMRIDatabase/).

## Acknowledgements

LMD is supported by the Biotechnology and Biological Sciences Research Council (grant number BB/T008709/1). PP is supported by the Wellcome Trust (212979/Z/18/Z).

## Code, Data, and Materials Availability

The data supporting the results of this paper can be made available upon reasonable request to the corresponding author through a formal data sharing and project affiliation agreement.

